## Supplementary material for "plasma: Partial LeAst Squares for Multiomics Analysis": Suppplemental Material

#### Supplemental Results

##### Supplemental Figure S1

**Figure S1** shows that there is no significant difference in survival between the training and test sets formed by splitting the STAD TCGA data.

##### Supplemental Figure S2

**Figure S2** presents the performance of individual omics models on the STAD test data.

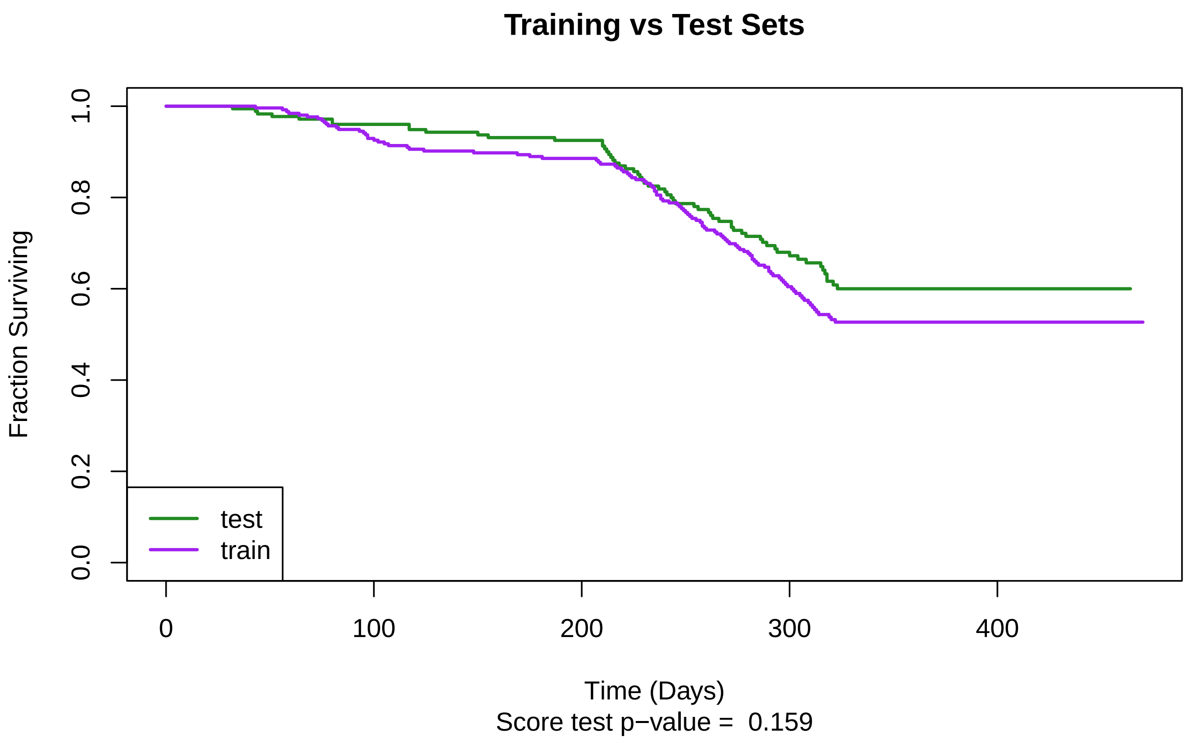


Figure 1: Kaplan-Meier plot of overall survival for the training and test sets.

#### Supplemental Figure S3

**Figure S3** is a heatmap equivalent to **Figure 7** in the main text, using correlation as the similarity measure to drive clustering. The vertical dendrogram makes it clear that the biological components associated with the "final" joint model are most strongly correlated with those associated to "all genes" used across individual components. The color scheme also highlights the fact that biological processes in the center part of the plot tend to be strongly associated with all components used in producing the final prognostic model.

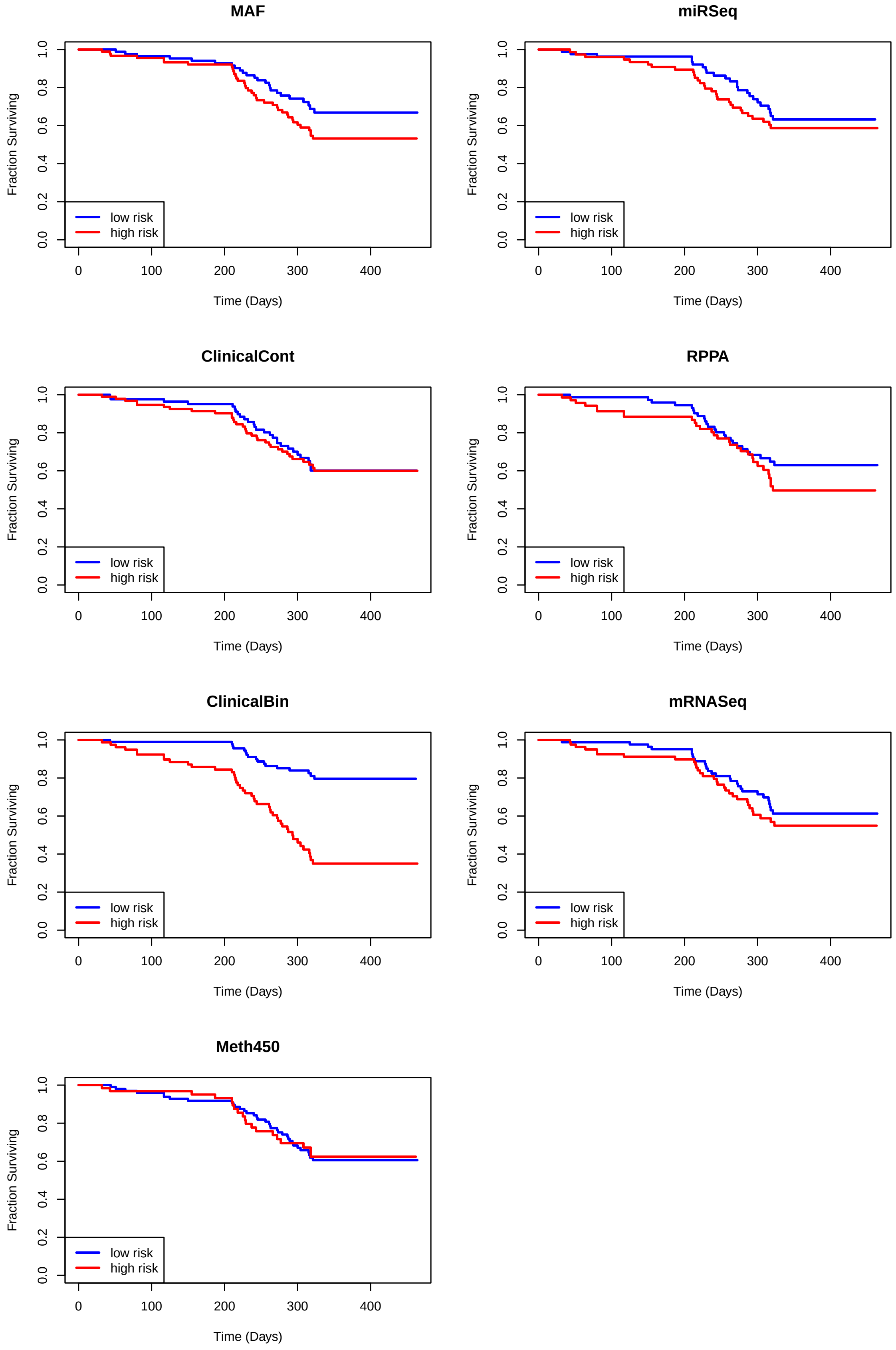


Figure 2: Kaplan-Meier plots of overall survival on the STAD test set from separate PLS Cox omics models.

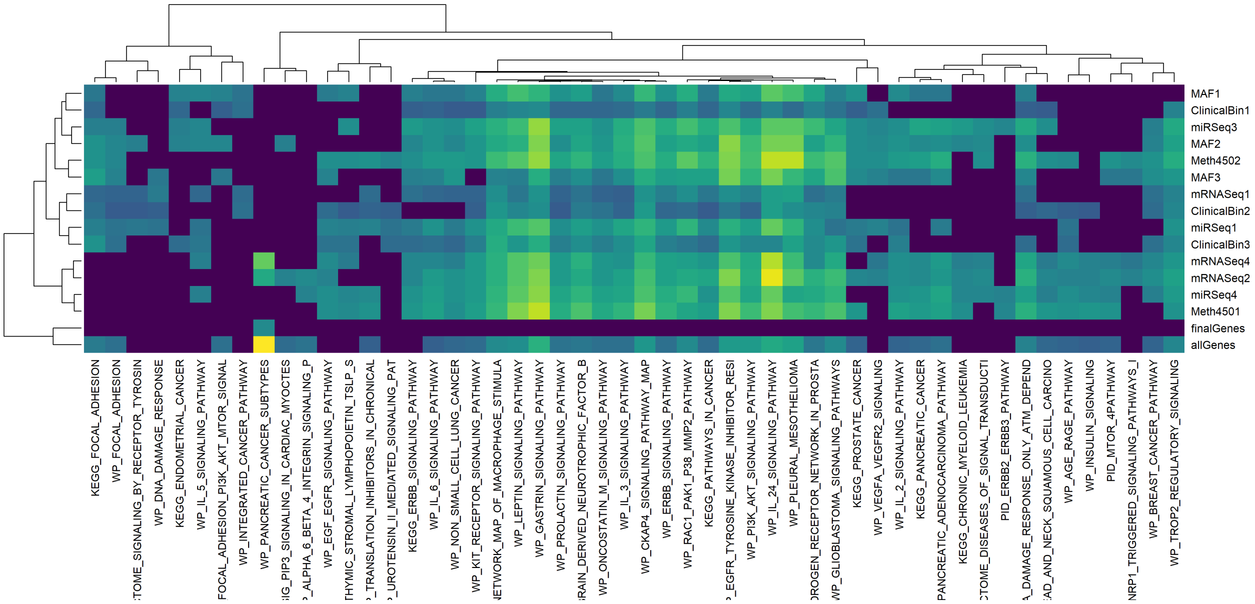


Figure 3: Components clustered based on biological process annotation p-values.

#### Supplemental Figure S4

**Figures S4 - S7** are bubble plots similar to **Figure 7** in the main text, showing the association of the components with different kinds of gene sets.

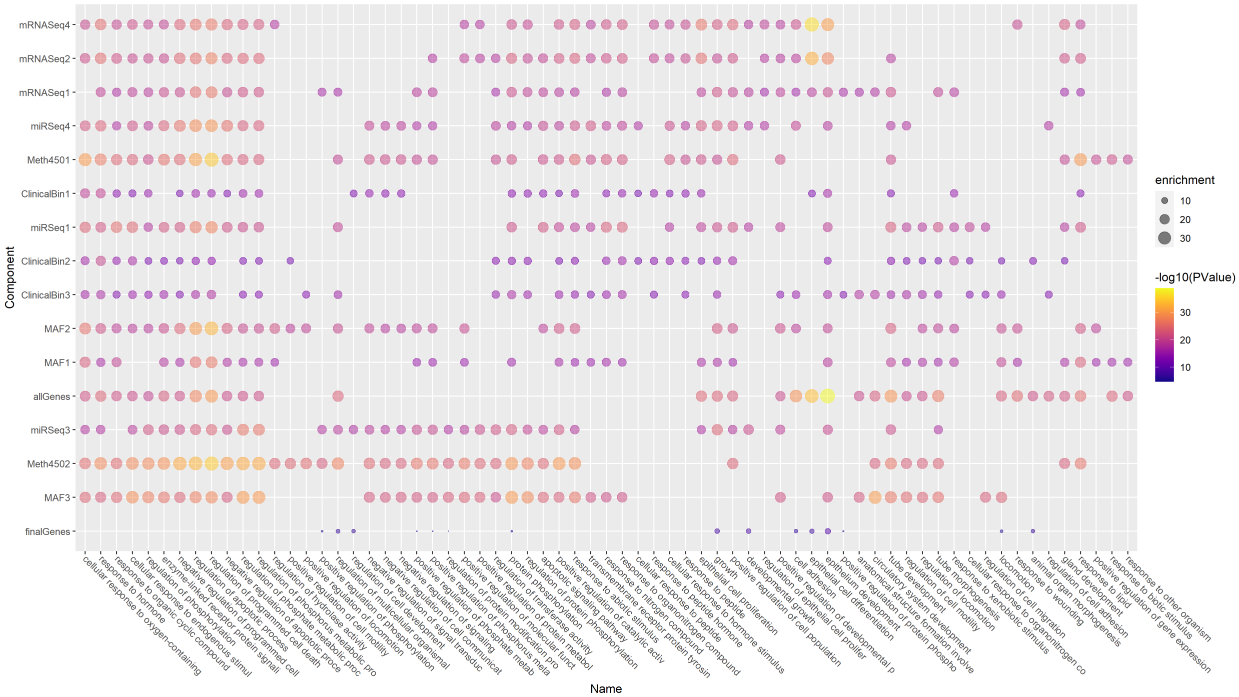


Figure 4: Components clustered based on biological process annotation p-values.

#### Supplemental Figure S5

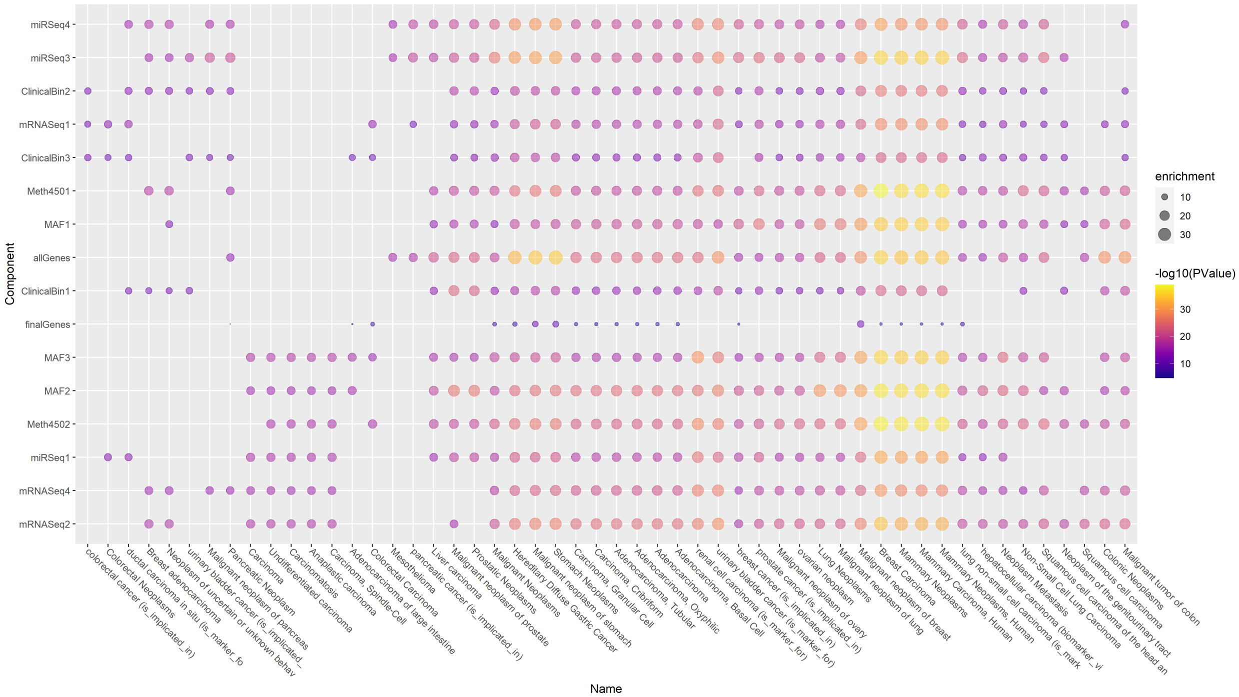


Figure 5: Components clustered based on disease annotation p-values.

#### Supplemental Figure S6

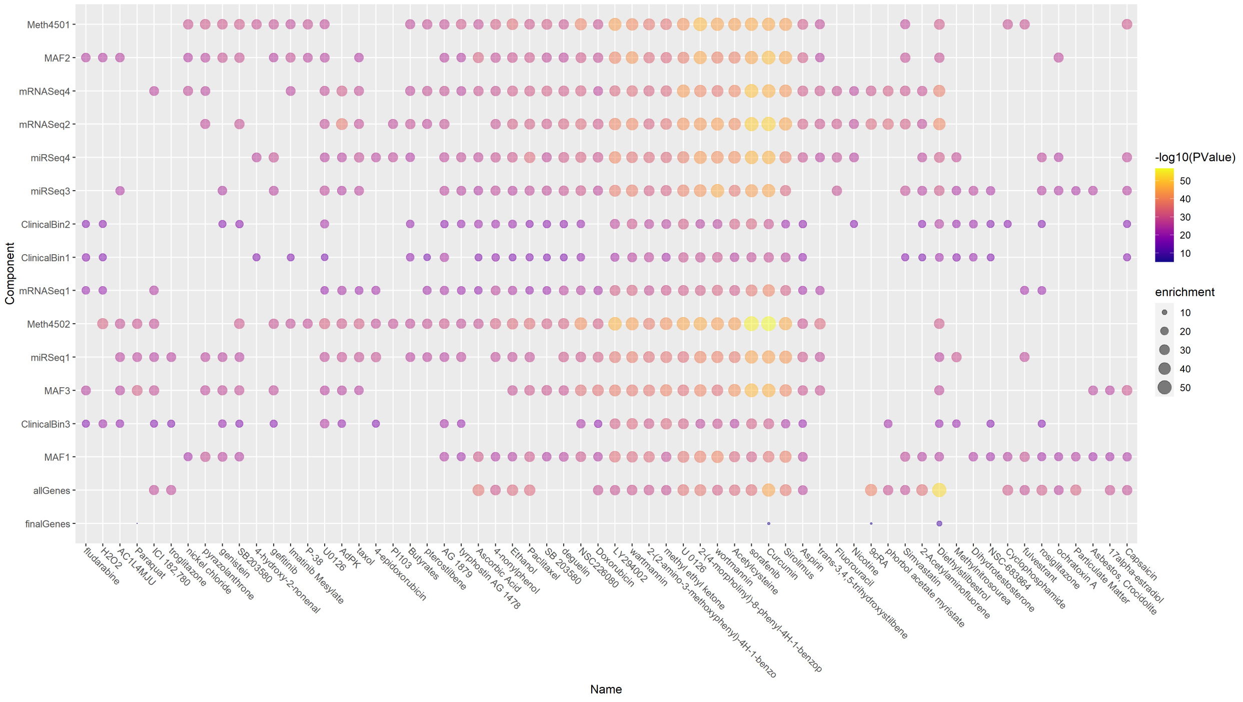


Figure 6: Components clustered based on drug annotation p-values.

#### Supplemental Figure S7

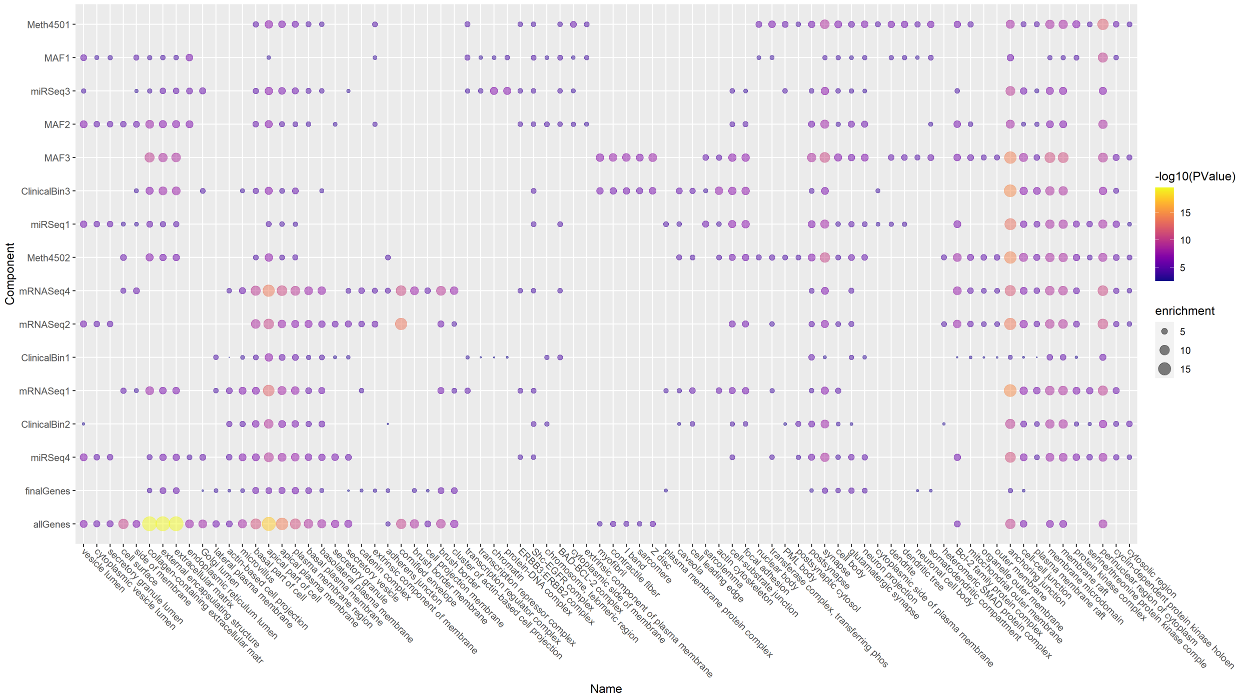


Figure 7: Components clustered based on cellular component annotation p-values.

### MOFA and Survival

Here, we run Multi-Omics Factor Analysis (MOFA) to illustrate the differences between a supervised analysis (as provided by our `plasma` package) and a mainly unsupervised analysis of multi-omics data to predict time-to-event outcomes. By the latter, we specifically mean

1. Running MOFA to detect latent factors across multiple omics data sets.
2. Testing the latent factors on new data to see if they are at all associated with the time-to-event outcome that we want to be able to predict.

We load the same version of the data, already split into training and test sets, that we used for our `plasma` analysis in the main text. Then we create the initial MOFA object.

#### Supplemental Figure S8

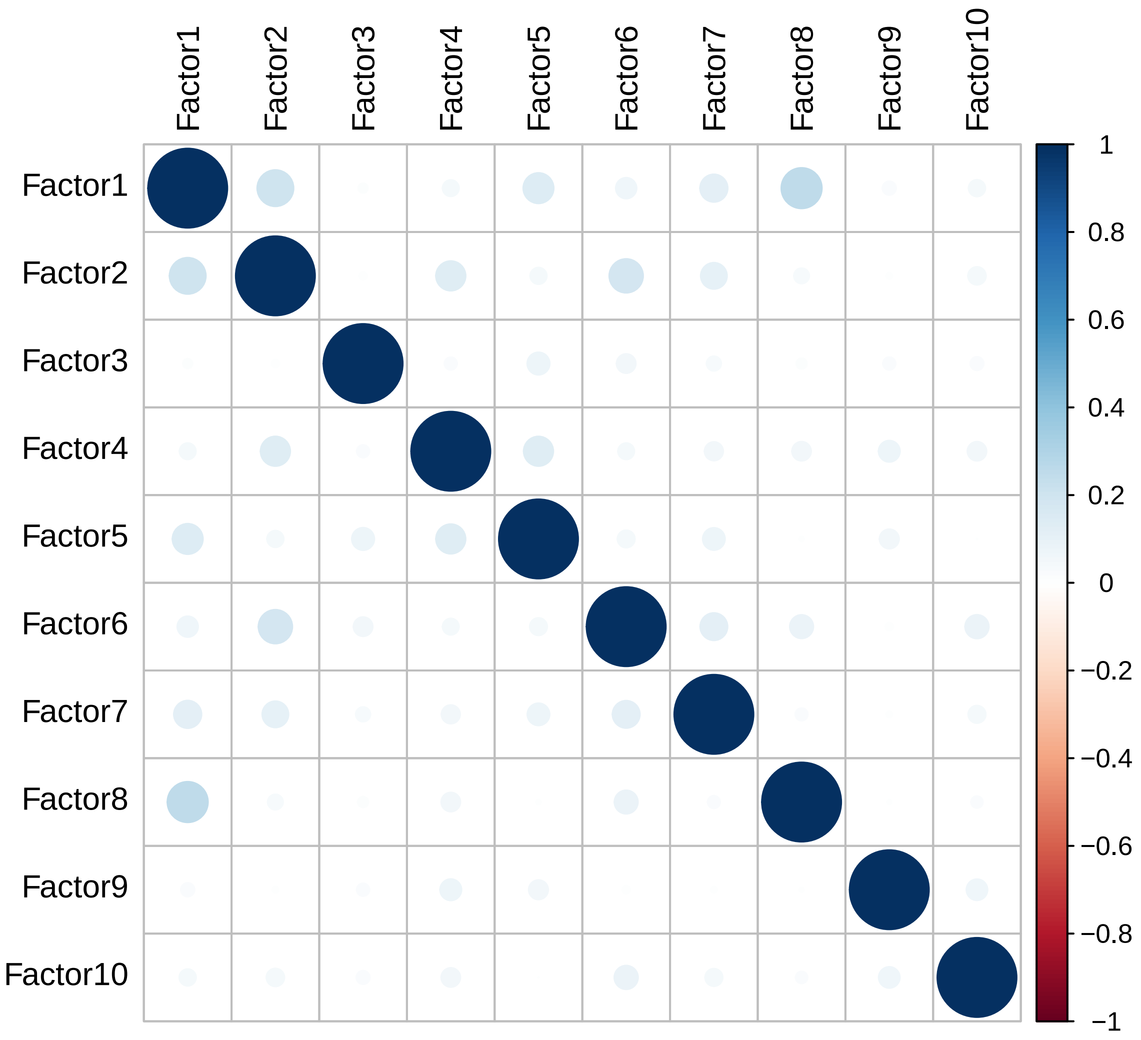


Figure 8: Correlation between MOFA latent factors.

#### Supplemental Figure S9

#### Training: Relationship to Overall Survival

We want to see whether any of the latent factors are associated with overall survival. We start by fitting separate univariate models for each of the latent factors. The results of these univariate models suggest that only Factors 2, 10, and 4 are even marginally useful for prediction (**Table T1**).

Table 1: Table T1: Details of the univariate Cox Proportional Hazards models for overall survival (OS) using MOFA factors as predictors (Coef = log hazard ratio, HR = hazard ratio, score = log-rank test statistic).

| | coef | HR | score | pvalue |
|---|---|---|---|---|
| Factor1 | 0.0460390 | 1.0471153 | 0.9122469 | 0.3395189 |
| Factor2 | 0.0969772 | 1.1018352 | 2.5846875 | 0.1079017 |
| Factor3 | 0.0217074 | 1.0219448 | 0.0750634 | 0.7841023 |
| Factor4 | -0.1168017 | 0.8897616 | 1.4352500 | 0.2309095 |
| Factor5 | 0.0409005 | 1.0417485 | 0.2412707 | 0.6232898 |
| Factor6 | 0.0089701 | 1.0090104 | 0.0086321 | 0.9259759 |
| Factor7 | 0.0862323 | 1.0900596 | 0.8023007 | 0.3704064 |
| Factor8 | -0.0190968 | 0.9810844 | 0.0446093 | 0.8327241 |
| Factor9 | 0.0647337 | 1.0668749 | 0.4330933 | 0.5104750 |
| Factor10 | 0.0948066 | 1.0994462 | 1.5297208 | 0.2161543 |

Next, we construct a multivariate model combining all of the factors as possible predictors of overall survival. The p-values associated with individual factors in the multivariate model again suggest that, at best, only Factors 2 and 4 might be useful (**Table T2**).

Table 2: Table T2: Details of the multivariate Cox Proportional Hazards model for overall survival (OS) using all MOFA factors as predictors (coef = log hazard ratio, exp(coef = hazard ratio. se = standard error).

| | coef | exp(coef) | se(coef) | z | Pr(>\|z\|) |
|---|---|---|---|---|---|
| Factor1 | 0.0177368 | 1.0178950 | 0.0548793 | 0.3231966 | 0.7465463 |
| Factor2 | 0.1140809 | 1.1208428 | 0.0669179 | 1.7047875 | 0.0882341 |
| Factor3 | 0.0433160 | 1.0442678 | 0.0801522 | 0.5404217 | 0.5889062 |
| Factor4 | -0.1488849 | 0.8616683 | 0.0992794 | -1.4996561 | 0.1337035 |
| Factor5 | 0.0479219 | 1.0490888 | 0.0842644 | 0.5687093 | 0.5695534 |
| Factor6 | -0.0391104 | 0.9616445 | 0.1038272 | -0.3766879 | 0.7064055 |
| Factor7 | 0.0727848 | 1.0754990 | 0.1050473 | 0.6928765 | 0.4883870 |
| Factor8 | -0.0118319 | 0.9882379 | 0.0915264 | -0.1292726 | 0.8971419 |
| Factor9 | 0.0749702 | 1.0778521 | 0.0973015 | 0.7704946 | 0.4410066 |
| Factor10 | 0.1065461 | 1.1124292 | 0.0755382 | 1.4104933 | 0.1583941 |

Finally, we use the Akaike Information Criterion (AIC) to formally select factors to retain in the final model. This process confirms our earlier suspicions that only Factors 2 and 4 make useful contributions to the model (**Table T3**). Starting with the predicted risk scores from the AIC model, we cut the scores at 0 to create "low" and "high" risk cohorts. The resulting Kaplan-Meier plot is shown in **Figure S10**. The model has a not-quite significant separation between the two risk groups (p = 0.06) in the training set.

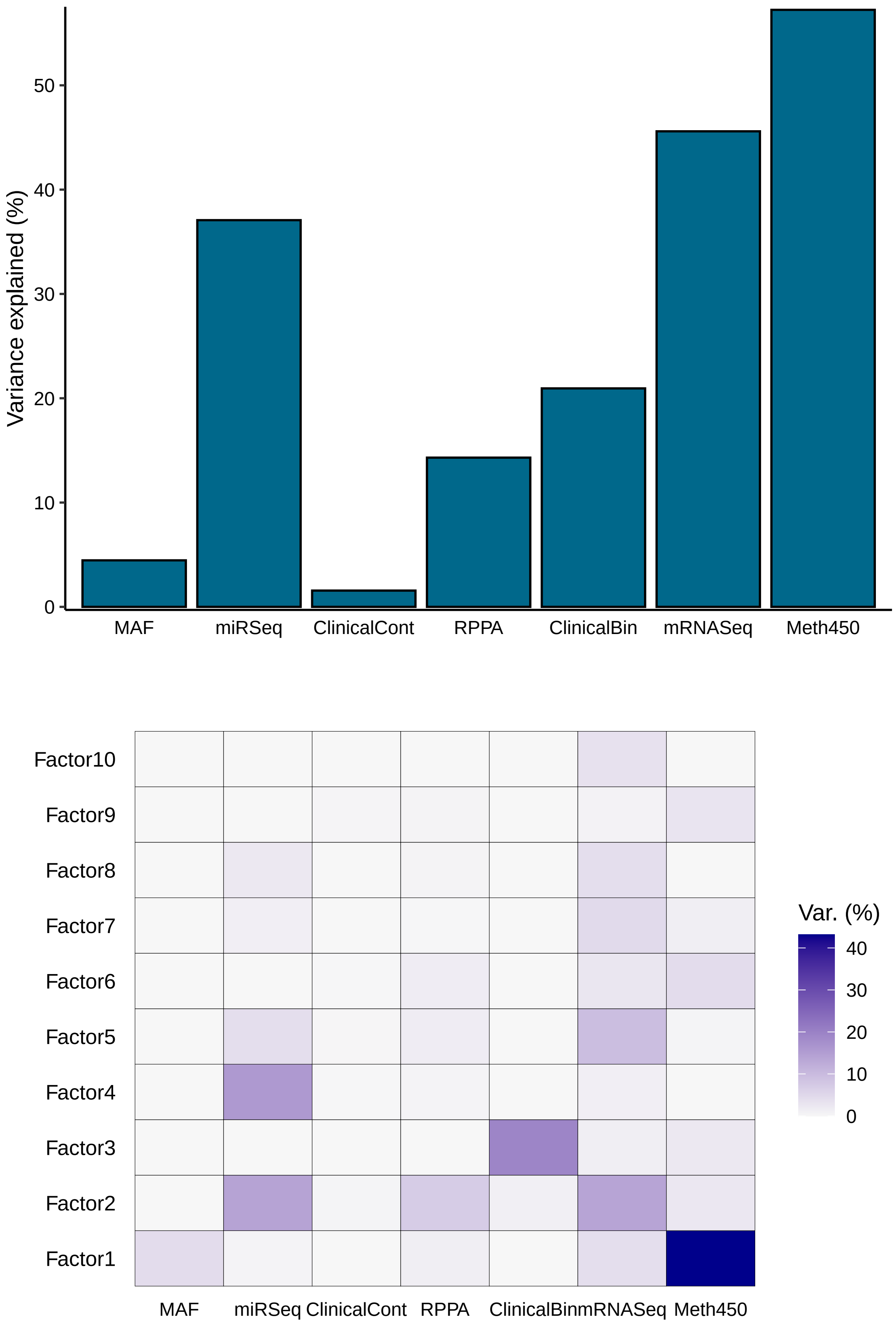


Figure 9: Amount of variance explained by latent factors in omics data sets.

Table 3: Table T3: Details of the multivariate Cox Proportional Hazards model for overall survival (OS) with variables selected using stepwise regression with AIC.

| | coef | exp(coef) | se(coef) | z | Pr(>\|z\|) |
|---|---|---|---|---|---|
| Factor2 | 0.1119490 | 1.118456 | 0.0621418 | 1.801508 | 0.0716228 |
| Factor4 | -0.1402108 | 0.869175 | 0.0962488 | -1.456753 | 0.1451846 |

**Supplemental Figure S10**

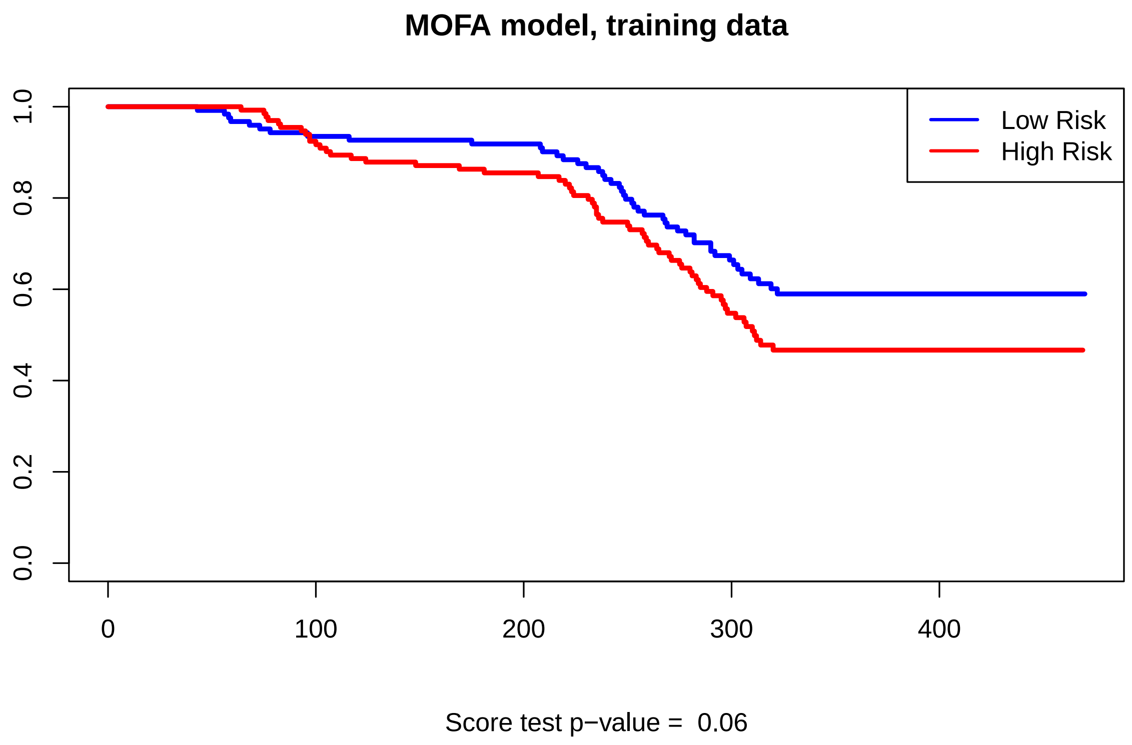


Figure 10: Kaplan-Meier plot based on risk predicted by MOFA latent factors.

#### Approximating the MOFA Computations

The hard part is to reverse-engineer the MOFA model so we can use its decompositions to make predictions on the test data. The statistical model underlying MOFA is that the matrix $Z$ of latent factors should simultaneously satisfy the equations $X_i = W_i Z$ for each omics data matrix $X_i$ and for appropriate weight matrices $W_i$. In principle, this means that we should (at least approximately) be able to recompute $Z$ from the original data matrices and the known weight matrices.

First, we illustrate two ways to recover the latent factor matrix $Z$.

```
Z0 <- get_factors(mofaSTADedit)[[1]]
Z <- mofaSTADedit@expectations$Z[[1]]
identical(Z0, Z) # TRUE
```

```
## [1] TRUE
```

Next, we illustrate two ways to recover the weight matrices, $W_i$.

```r
weightmats <- get_weights(mofaSTADedit)
identical(weightmats, mofaSTADedit@expectations$W) # TRUE
```

```
## [1] TRUE
```

```r
Woo <- do.call(rbind, weightmats)
```

The last line in the previous code chunk bundles the weight matrices together into a single matrix $W$.

Because MOFA internally centers the data in continuous input matrices, we have to do the same thing to the input training data.

So, we bundle the (centered) input data matrices into a single matrix, $X$. We also replace all of the missing data (which arises as complete rows when a patient has not been included in an assay) with zeroes. Then we solve for $Z$ by using a QR-decomposition of the full weight matrix.

##### Supplemental Figure S11

In **Figure S11**, we show that the approximation that we have constructed for $Z$ still yields a model that separates the training data into low- and high-risk patients. Because this only approximates the actual MOFA latent factors, this model is even less significant (p=0.11) then the previous one.

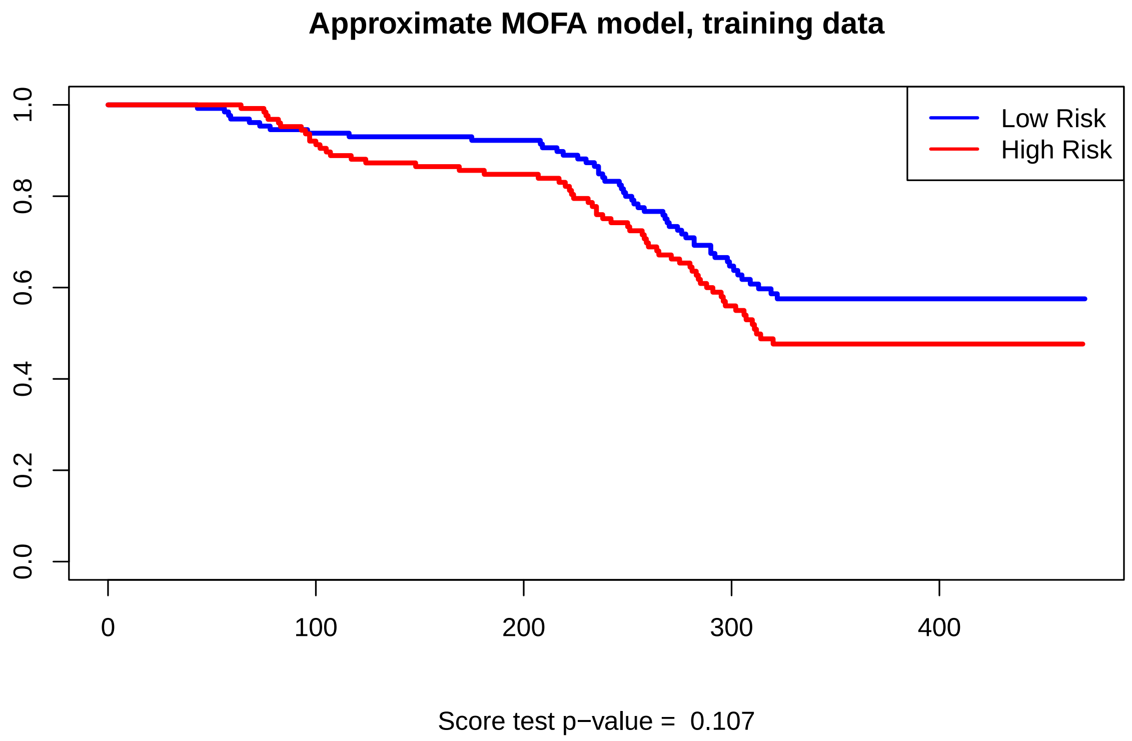


#### Test Data

Now we can use the same weight matrices that we learned on the training data to project the independent test data into the latent factor space.

Then we split the predicted scores from the AIC model applied to the test data at the same cutoff (i.e., zero) that we used for training.

##### Supplemental Figure S12

Surprisingly (since the model was not significant on the trainng set, it does seem to do a passable job on the independent test data, with p = 0.016 (**Figure S12**). This result is soewhat disconcerting. Any sort of learned model isn't supposed to perform *better* on the test set than it does on the training set. We suspect that we just got "lucky".

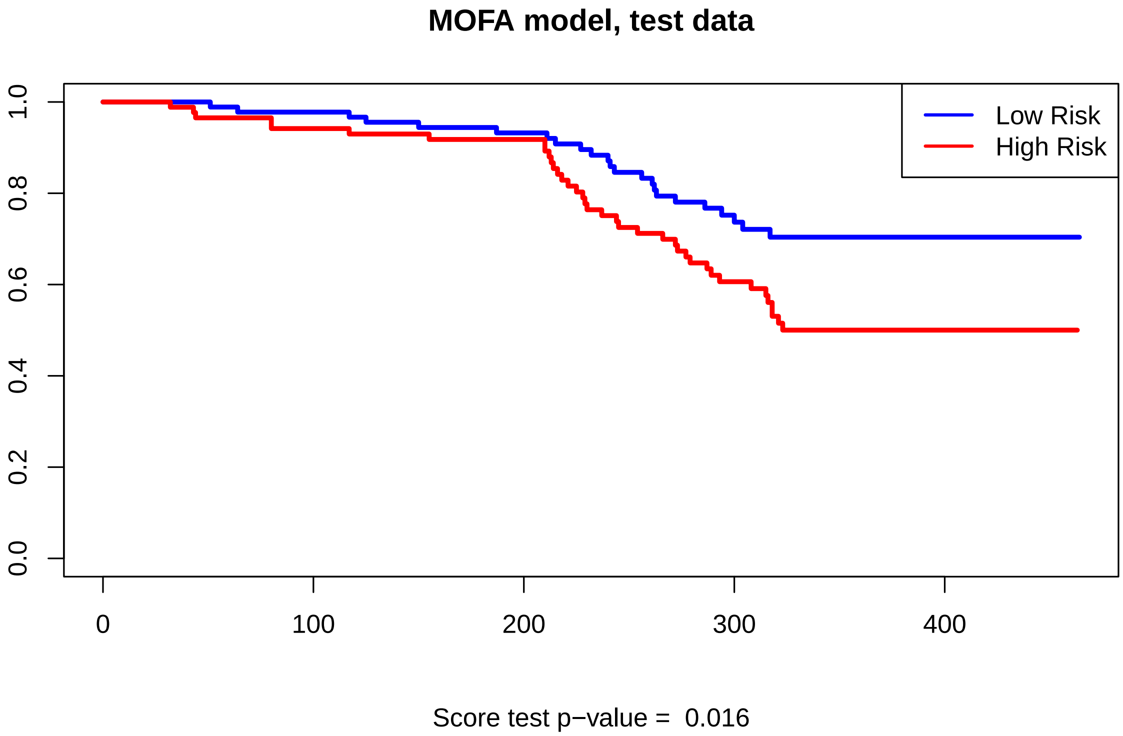


Figure 11: Kaplan-Meier plot of MOFA model on test data.

#### Independent Validation with ESCA Adenocarcinoma Data

Finally, we apply the same model that we we learned from the STAD training data to the adenocarcinoma subset of the esophageal cancer (ESCA) data from TCGA.

##### Supplemental Figure S13

**Figure S13** shows that the model is completely unable to predict prognosis in the ESCA adenocarcinoma data set (p = 0.237)

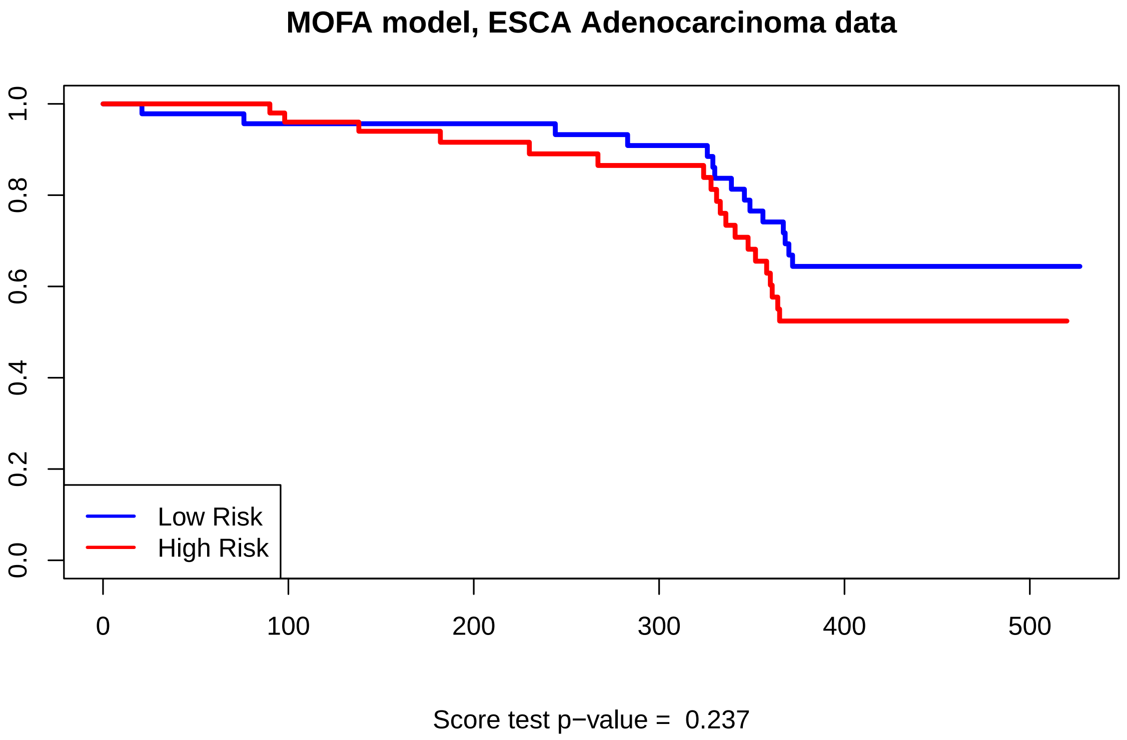


Figure 12: Kaplan-Meier plot of MOA model on the ESCA adenocarcinoma validation data.
